# Torsion-Induced Traumatic Optic Neuropathy (TITON): A Physiologically Relevant Animal Model of Traumatic Optic Neuropathy

**DOI:** 10.1101/2024.10.08.617086

**Authors:** Annie K. Ryan, Brooke I. Asemota, Tyler Heisler-Taylor, Claire Mello, Luis Rodriguez, William E. Sponsel, Julie Racine, Tonia S. Rex, Randolph D. Glickman, Matthew A. Reilly

## Abstract

Traumatic optic neuropathy (TON) is a common cause of irreversible blindness following head injury. TON is characterized by axon damage in the optic nerve followed by retinal ganglion cell death in the days and weeks following injury. At present, no therapeutic or surgical approach has been found to offer any benefit beyond observation alone. This is due in part to the lack of translational animal models suitable for understanding mechanisms and evaluating candidate treatments. In this study, we developed a rat model of TON in which the eye is rapidly rotated, inflicting mechanical stress on the optic nerve and leading to significant visual deficits. These functional deficits were thoroughly characterized up to one week after injury using electrophysiology and immunohistochemistry. The photopic negative response (PhNR) of the light adapted full field electroretinogram (LA ffERG) was significantly altered following injury. This correlated with increased biomarkers of retinal stress, axon disruption, and cell death. Together, this evidence suggests the utility of our model for mimicking clinically relevant TON and that the PhNR may be an early diagnostic for TON. Future studies will utilize this animal model for evaluation of candidate treatments.

## Introduction

Traumatic optic neuropathy (TON) is closely associated with traumatic brain injury (TBI), and results in irreversible blindness (1). TON occurs in up to 5% of all closed head injuries and 2.5% of maxillofacial and mid-face traumas (2–4). This results in upwards of 75,000 people per year developing irreversible vision loss as a result of TON. Ocular injuries are the 4^th^ most common battlefield injury and, in the US, make up around 2.5 million emergency department visits per year (5,6).

Intervention relies primarily on high dose corticosteroid administration and/or surgical decompression; however, both have been observed to have limited success (2,3,7,8). In addition, treatment with both corticosteroids and surgical decompression have been called into question due to the increasing evidence of a higher risk of morbidity when administered in TBI patients (3,8–10). In this study, 21% of TBI participants who were administered corticosteroids died at 2 weeks whereas in the placebo group only 18% died within this timeframe (10). The authors concluded, corticosteroids should not be utilized to treat TBI patients (10). Due to the common coexistence of TBI and TON, corticosteroid administration has been minimized, resulting in an urgent need for therapeutic intervention before vision loss occurs. The first step towards this goal is development of a physiologically-relevant animal model in which to determine the cellular injury pathway and find a diagnostic criterion capable of locating visual dysfunction before significant cell loss occurs.

In general, TON is characterized by a loss of vision associated with retinal ganglion (RGC) death and axonal loss of the optic nerve; however, the exact mechanism by which the visual system is affected is not fully understood (1,11,12). Translationally relevant animal models are limited with most studies relying on optic nerve crush or transection. These models focus on a direct injury inducement, but do not address the more common indirect causes of injury (13). One indirect TON small animal model utilizes air blast trauma to develop a repeatable injury (14). This model serves to mimic the improvised explosive device blast many military personnel experience; however, this model does not mimic the torsional shearing that is sometimes observed in other forms of coup-contrecoup TBI. In our model we aim to develop a torsional indirect traumatic optic neuropathy (TITON) event.

Our injury mechanism more closely aligns with closed head TBIs as it induces a torsional shearing event similar to what the optic nerves experience during the coup-countercoup motion of the brain during the acceleration and deceleration event of TBI (8,15,16). After coup-contrecoup motion, deformational injuries to the optic nerve such as bending, shearing, and compression have been reported (15,16). As the optic nerve experiences biomechanical stress, the retina can experience injury events as well. RGCs are the cell type on the anterior portion of the retina, with axons that extend posteriorly and bundle together to form the optic nerve. Thus, since the optic nerve is largely comprised of RGC axons, this injury event may translate to the retina. The direct method in which vision loss occurs and which cell types are initially afflicted is debated. Determination of the cells in the retina and optic nerve that are injured is crucial to determining diagnostic and treatment protocols. We therefore employed light adapted full field electroretinograms (LA ffERGs) to find early diagnostic protocols and to determine the cell types afflicted after TITON.

In this study we developed a translationally relevant small animal model of TON via a torsional rotational injury event, similar to closed head TBI. This TITON model was thoroughly characterized using ffERG. Based on these observations, we also propose the LA ffERG as a means of early diagnostic screening for TON.

## Materials and Methods

The overall experimental design was comprised of two phases. First, a pilot study was conducted to estimate the ocular rotation required to achieve a low level of irreversible functional loss. Second, a more extensive characterization of the low-level injury was undertaken to evaluate potential diagnostic biomarkers.

### Animals

For the pilot study, 27 adult female Sprague Dawley rats (350-400g; Charles River Laboratories International, Inc., Wilmington, MA) were utilized to estimate amplitude, velocity, and degree of rotation required to achieve a low-level, irreversible injury. For the current study, 44 ∼200g male Sprague Dawley rats were used (Charles River). All experiments were conducted in accordance with the ARVO Statement for the Use of Animals in Ophthalmic and Visual Research and institutionally approved protocols. Sham animals in this study are comprised of our uninjured cohort, which have undergone sham operations and maintain the same number of anesthetic events. Contralateral eyes of animals that received the injury event were used as a second level control. The contralateral eyes of these animals were not directly injured via our TITON event.

### Injury Induction

A robot was constructed (**Fig. 1**) to allow the application of rapid, coaxial rotation to the eyes of a rat. The foundation of the instrument was a rat stereotaxic (Mouse Specific Stereotaxic Base; Leica Microsystems, Inc., Richmond, IL) that allowed complete immobilization of the rat’s head and accurate positioning of the actuator relative to the superior-inferior axis of rotation of the left eye. The actuator (BLM-N23; Galil Motion Control, Rocklin, CA) was fitted with a quadrature encoder (4000 counts/revolution) yielding an angular resolution of 0.09 degrees. The actuator was controlled using a programmable motion controller (DMC-30012; Galil Motion Control) that allowed prescription of maximum angular displacement, velocity, and acceleration.

**Fig. 1:**
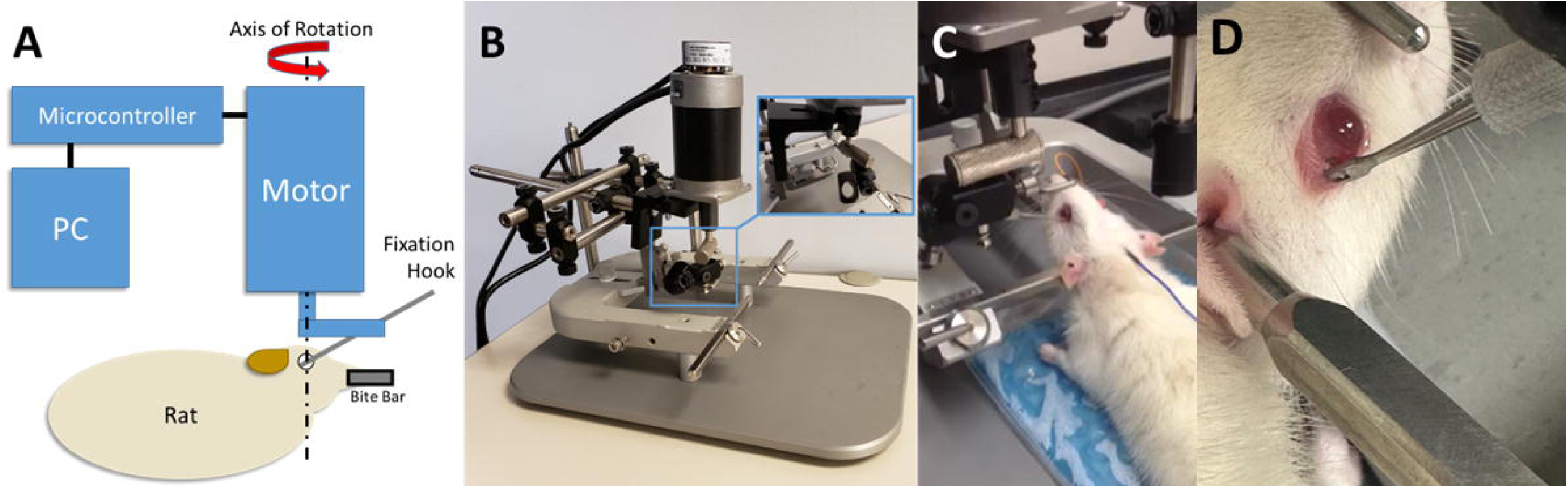
Schematic and photographs of the TITON apparatus. (A) The instrument allows independent variation of the amplitude, rate, and direction of rotation to either eye of the rat via a custom user interface. (B) A motor was rigidly mounted on a stereotaxic unit to allow precise positioning of the motor relative to the eye’s axis of rotation while immobilizing the animal’s head to ensure accurate rotation. (C) Rat in stereotaxic with fixation hook attached to left eye. (D) Rat in stereotaxic with fixation hook post TITON inducement.

Prior to induction of the traumatic event, an initial calibration of the robot apparatus was conducted using the control software (GalilTools; Galil Motion Control) at the desired parametric rate and amplitude of rotation to ensure proper connectivity of the motor and ensure that the eye and rig were able to achieve the desired range of motion. A scleral fixation hook (47-2650 Guthrie Fixation Hook; Sklar Instruments, West Chester, PA) was used to rigidly connect the axis to the temporal extraocular musculature (Fig. 1C). For the pilot study the left eyes of animals were utilized while in the current study the right eye was subjected to rotations up to 47 degrees at angular velocities up to 3320 degrees/s. Peak amplitude and velocity were then recorded.

### Identification of Injury Thresholds

Flash visual evoked potentials (fVEPs) were recorded before, immediately after injury, and again seven days post injury to quantify the extent of injury. The following experimental breakdown of animal cohorts were utilized: negative controls (no eye rotation) n=3; positive controls (normal saccade-level rotation) n=3; low n=4, medium n=8, and high n=10.The animal was anesthetized throughout visual testing and the trauma event with ketamine (80 mg/kg) and xylazine (8 mg/kg), which was administered intraperitoneally. Following anesthesia, a thin layer of optical ointment (neomycin and polymyxin B sulfates, and bacitracin zinc ophthalmic ointment; Akorn, Inc., Lake Forest, IL) was applied to the eye to prevent drying and infection.

fVEPs utilized a ∼10μs flash was presented by an external stimulator (Grass PS22D Photic Stimulator, Grass RPS107 Regulated Power Supply; Grass Instrument Co., Quincy, MA). These light flashes induced fVEPs in the corresponding visual cortex of the brain. The fVEPs were sampled as an average of 64 signal data points and were acquired using sub-dermal electrodes (Low Profile Needle Electrode for EEG; Chalgren Enterprises, Inc., Gilroy, CA; TDS 2014C Oscilloscope, Tektronix Inc., Beaverton, OR; IGMEB-NUM25 Mini Electrode Board; Grass Inst. Co., Quincy, MA; 12B-8-23 Neurodata Acquisition System; Grass Inst. Co., Quincy, MA) that were placed to maximize the signal-to-noise ratio.(17) Three data sets were collected as follows. One minute of dark adaptation preceded fVEP followed by one minute of dark adaptation prior to collection of a control waveform recorded in an identical way except that the eye was not exposed to the flash stimulus to ensure the recorded activity was a direct result of the external stimulus. Three such data sets were recorded at each time point immediately before, immediately after, and one week after trauma.

After the fVEPs had been taken following the traumatic event, the rat was given 2.1 mg/kg yohimbine (Yobine®; Lloyd Laboratories, Shenandoah, IA) to reverse the initial anesthesia. One week after the initial session, the rat was anesthetized as above and another set of fVEPs were acquired using the previously mentioned protocol.

Changes in fVEP amplitude and latency at N1 and P1 were normalized against baseline readings to account for inter-animal variability. These metrics were computed for fVEPs immediately after injury (D0) and seven days after injury (D7).

Negative controls were placed in the apparatus and removed without applying any ocular rotation. Positive controls were subjected to saccade-level ocular rotation. An example of the angular position and velocity from one experiment is shown in **Fig. 2 A & B, respectively**. Signal – control fVEP waveforms were used to measure the difference in amplitude and latency from N1 to P1 (**Fig. 2C**). A threshold-based statistical model was used to estimate the angular rotation required to achieve a functional deficit immediately after injury as well as 7 days after injury.

**Fig. 2:**
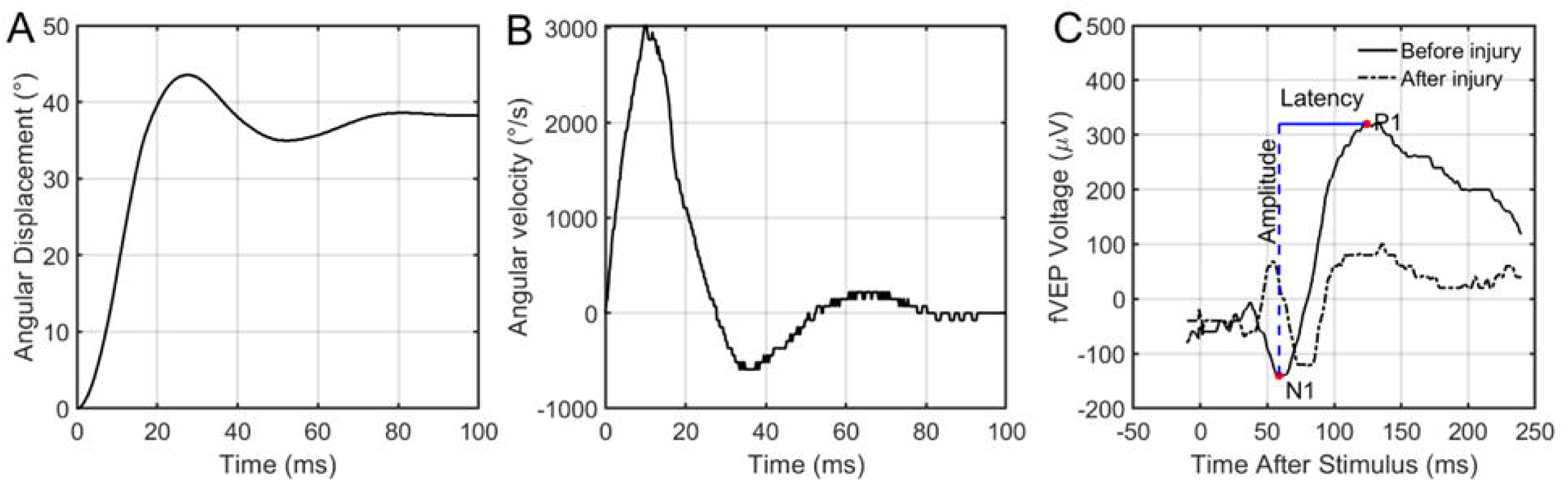
Representative ocular rotation and fVEP data. Representative example of the rapid rotation applied to the specified (A) angular displacement (45 degrees) and (B) peak velocity (3000 degrees/second) to the left eye of a rat. (C) Sample fVEP before and immediately after ocular rotation shown in A and B. Overlaid is a schematic data extraction procedure for an fVEP waveform. The waveform itself was generated by collecting and averaging numerous signals following flash exposure, then subtracting an averaged signal collected in the same way without the flash.

### Functional Characterization of the Injury

Prior to testing, rats (n=39 total; n=31 injured; n=8 sham) were anesthetized with an IP injection of a mixture of ketamine (West-Ward; Eatontown, NJ) and xylazine (Akorn Inc.; Lake Forest, IL) (90mg/kgketamine and 10mg/kg xylazine). Once the pedal withdrawal reflex was absent, the eyes were dilated with 1% Tropicamide Ophthalmic Solution (Akorn Inc.). In preparation for functional characterization, one platinum subdermal needle electrode (Natus Manufacturing Limited; Gort, CO. Galway Ireland) was placed in the tail, another subdermal needle electrode (Natus) was placed in the center of the head, one 6mm gold surface cup electrode (Natus) was placed in the mouth and finally, after the eyes were numbed with 0.5% Tetracaine Hydrochloride Ophthalmic Solution (Bausch & Lomb Inc.; Tampa, FL) and lubricated with Systane gel (Alcon Laboratories Inc.; Fort Worth, Texas) or GentleTears gel (Alcon), the Celeris electrodes (Diagnosys LLC; Lowell, MA) were placed perpendicularly on the surface of each eye slightly touching the cornea. To record the full field electroretinograms and the flash visual evoked potentials, the Celeris system (Diagnosys LLC; Lowell, MA) was used. During the entire testing period, the internal temperature of the rats was kept at 37°C.

LA ffERGs, oscillatory potentials (OPs) and fVEPs were obtained against a background of 30 cd/m^2^ using a white flash (flash intensity: 200 cd.s/m^2^) with an interstimulus interval (ISI) of 1000ms. An average of 50 sweeps were collected. The light-adapted flicker ERGs were also recorded. A white flash (flash intensity: 200 cd.s.m^2^) was used. A total of 200 sweeps were collected with a frequency of 20 Hz. Finally, the photopic negative response (PhNR) were also recorded and used a white stimulus (flash intensity: 200 cd.s/m^2^) against a green background (40 cd/m^2^). An average of 100 sweeps were collected.

A-waves were measured from baseline to the first negative trough. The b-waves were measured from the a-wave peak to the highest peak of the response. Flicker ERG and OPs were measured from preceding trough to peak. OPs were calculated utilizing the sum of the OPs (SOP) and only OPs that occur prior to the peak of the b-wave were utilized (SOP= OP1-2 + OP3 + OP4 + OP5). PhNR amplitudes were measured from baseline to the trough following the b-wave. Peak times were measured utilizing the time of the flash onset to respective peaks.

Visual electrophysiology testing was performed at baseline (no more than 7 days before injury event) (n=31) and at 1 (n=26) and 7 (n=14) days (D1 and D7, respectively) after injury. The n values reflect the number of animals receiving ERGs at each time point.

Five D1 and one D7 ERGs were excluded due to insufficient data quality from excessive 60 cycle noise. Sham animals (n=8) received the same schedule of ERG testing with both D1 (n=7) and D7 (n=5) endpoints. One sham D1 ERG was excluded due to the above reasoning. At the end of the testing period, the animals were injected with an IP dose of Antipamizole (Modern Veterinary Therapeutics LLC; Miami, FL) (2mg/kg) to reverse the effect of xylazine and ease the recovery of the animals.

### Histology and Immunohistochemistry

Animals were humanely euthanized immediately following D1 or D7 ERG measurements using established and approved IACUC protocol as described below. Animals underwent CO_2_ asphyxiation followed by a lateral incision of the atlanto-occipital joint in preparation of tissue collection. Briefly, the skull was removed. The brain was reflected rostrally to expose the optic nerves. The nerves were then separated from the brain, and the brain was removed. The nerves and eyes were then dissected en bloc for immunohistochemistry analysis. For immunohistochemistry, rats were perfused with phosphate-buffered saline (PBS) followed by 4% paraformaldehyde, and the eye and optic nerve were dissected and incubated overnight at 4°C in 4% paraformaldehyde. The optic nerves and chiasm were dissected away from the brain and cryo-preserved in 20% sucrose in PBS for 2 hrs at 4°C followed by 30% sucrose in PBS overnight at 4°C. Tissues were embedded in OCT (Fisher Scientific, Waltham, MA) and longitudinally sectioned on a cryostat at 10 µm thickness. For immunohistochemistry, sections were incubated in PBS to remove the OCT and then incubated in 1:20 normal donkey serum in PBS plus Triton-X-100 (PBT) at room temperature for 2 hrs. Sections were then incubated with specific antibodies as follows: anti-β-tubulin (1:1000; MAB5564; Millipore, Burlington, MA) and anti-glial fibrillary acidic protein (GFAP; 1:50; Z0334; DAKO, Santa Clara, CA) in PBT overnight at 4°C, rinsed with PBS, and incubated in donkey anti-mouse Alexa 488 and donkey anti-rabbit Alexa 594 (1:200; Fisher Scientific) in PBT overnight at 4°C. Finally, the sections were rinsed, mounted in Vectashield plus DAPI (Vector laboratories, Burlingame CA). Tissue samples were imaged on a wide-field fluorescence microscope (Nikon Eclipse, Melville, NY) or confocal microscope (Nikon AXR) using consistent settings. The following n-values were used for each timepoint: sham retina and nerve (n=2), D7 retina (n=9), D7 nerve (n=3), D1 retina (n=3), D1 nerve (n=3).

A terminal deoxynucleotidyl transferase (dUTP) nick end labeling (TUNEL) assay was performed to assay cell death. Eyes with optic nerves attached were assayed. A 28-gauge needle was utilized at the temporal limbus of the eye to allow better perfusion of 4% PFA to the ocular interior. Eyes and nerves were then placed in 4% PFA and gently agitated for 2 hours before 2x 15-minute PBS wash. The samples were then transferred to 30% sucrose and incubated overnight at 4°C, then embedded in OCT (Fisher Scientific, Waltham, MA) and longitudinally sectioned on a cryostat at 10 µm thickness. Slides were then heated for 10 minutes on low heat. Once at room temperature, slides were soaked in an ABT solution for 2x 15min cycles with gentle shaking. Slides were then washed 2x with PBS for 10mins. TUNEL staining was completed following manufacturer’s instructions (TMR red kit, cat#: 12156792910, Roche) and with previously reported methods (18). After TUNEL kit application slides were washed 2x in PBS for 5 minutes. Hoechst nuclear stain was then applied to the slides for 10 minutes in a humidity chamber at room temperature. Slides were washed in PBS for 5 minutes before coverslips were mounted using 50% glycerol. Samples were then imaged with a Nikon Eclipse Ts2R fluorescent microscope with a panda sCMOS camera with NIS-Elements AR software (18). TUNEL positive (TUNEL+) cells counts were determined through MATLAB code quantification. The following n-values were used for each timepoint: sham retina and nerve (n=1), D7 retina and nerve (n=7), D1 retina and nerve (n=4).

### Statistical Analysis

For the Injury Threshold Estimation pilot study, statistical analysis was utilized to determine whether a significant correlation existed between the change in fVEP amplitude and/or latency and the velocity and/or amplitude of the TITON-induced trauma. A linear threshold model was used to determine if supra-saccade level rotation induced TON. The hypothesis for this analysis was that eye rotation at amplitudes and velocities exceeding that of normal saccades would induce TON. Therefore, correlation of the fVEP characteristics with the TITON-induced trauma would be dependent on the location of the threshold.

Specifically, a linear threshold model for the change in fVEP response *Δy* of the form

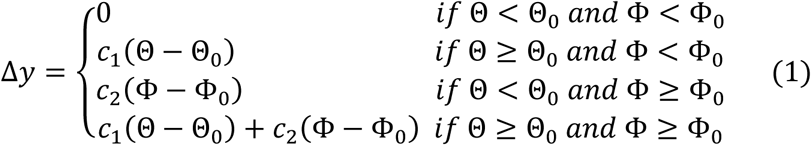

where *Θ* is the angular amplitude of the applied rotation, *Ω* is the angular velocity of the applied rotation, *Θ_0_* is the threshold value of angular amplitude required to achieve an irreversible injury, *Ω_0_* is the threshold value of angular velocity required to achieve an irreversible injury, and *c_1_* and *c_2_* are regression coefficients, was implemented in MATLAB r2015b (The Mathworks, Inc.; Natick, MA). Two changes in response were considered: the amplitude and P1-to-N1 latency of the fVEP signals (**Fig. 3**) collected immediately before and one week after ocular rotation.

**Fig. 3:**
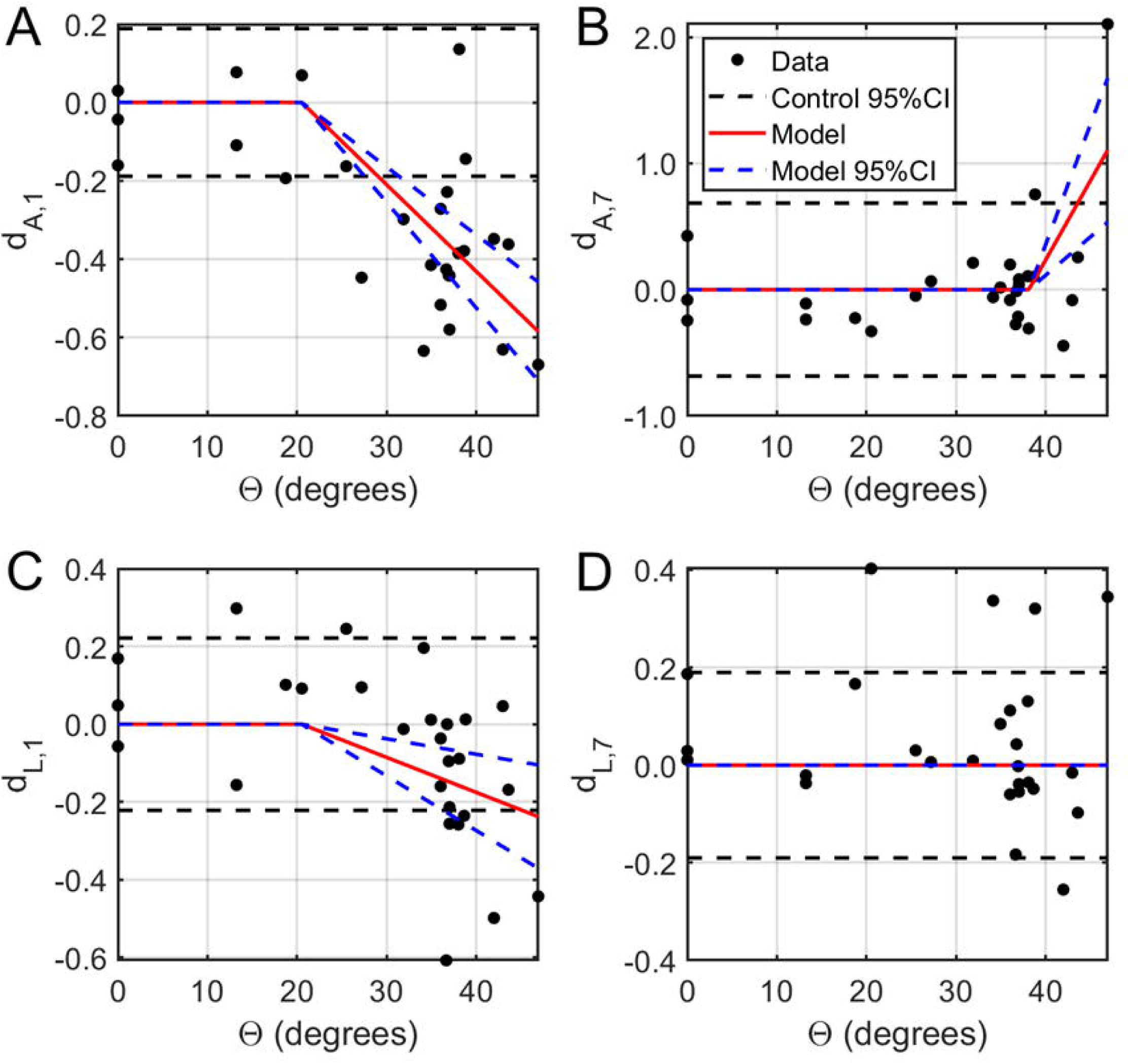
Estimation of injury thresholds from fVEP data. Normalized functional changes in fVEP amplitude (A, B) and latency (C, D) as a function of the angle of rotation *Θ*. Left: Both fVEP amplitude (A) and latency (C) were negatively correlated with the amplitude of ocular rotation above 20.5 degrees. Right: One week after ocular rotation, the fVEP amplitude increased with amplitude of ocular rotation above 38.1 degrees (B) while both fVEP amplitude and latency had higher variability. Subscripts 1 and 7 refer to the after-injury timepoint (i.e. either day 1 or day 7).Paired threshold values were postulated on a grid with *0° ≤ Θ_0_ ≤ 45°* and *0°/s ≤ Ω_0_ ≤ 3500°/s*. Linear regression was performed to estimate the regression coefficients and the F distribution was inverted to test for statistical significance. Contour plots showing lines of constant probability were generated in MATLAB. Thresholds were selected as the lowest pair of (*Θ_0_*, *Ω_0_*) values yielding a statistically significant, irreversible change in fVEP amplitude and latency at several significance levels (*α* = 0.05, 0.01, and 0.001). Threshold pairs were selected such that they were adequately supported by the available data. Thresholds were computed at three different significance levels because lower p values imply greater probability of finding irreversible injury and would therefore increase confidence in future animal studies; however, this likely also implies a more severe injury which may not be treatable and so these two factors must be balanced in designing future experiments.

Statistical Analysis for the Functional Characterization of the Injury study was performed using GraphPad Prism in which a two-wave ANOVA with a 95% confidence interval was performed between experimental groups. Tukey’s multiple comparisons tests were performed to further elucidate group interactions.

## Results

### Injury threshold estimation

Optic neuropraxia, defined as a reversible functional deficit in fVEP, was observed at insults with experimental extents of rotation between 20.5° - 38.1° (corresponding angular velocities 1475 - 3320°/s), which exceeded the maximal physiological saccadic parameters of 10° at 500°/s (**Fig. 3A,C**). Immediately after injury, fVEP amplitude decreased with faster peak times in a dose-dependent trend above 20.5° (effects of - 0.0221 and -0.0090 mV/°; p<0.0001 and 0.0011, respectively).

TON, defined as statistically significant changes from baseline measured one week after injury, was induced by rotations of larger than 38.1° with peak velocity greater than 3320°/s (**Fig. 3B,D**). Specifically, the final fVEP amplitude increased significantly relative to baseline above this threshold (0.1257 per degree; p=0.0005), while the variance in both amplitude and latency increased (p=0.0313 and 0.0317, respectively). The magnitude of the latency change was highly variable and did not exhibit any particular trend with respect to the angle of rotation (0.0113 per degree; p=0.8168).

### Functional characterization of injury

Significant alterations to the amplitude of the PhNR were observed following injury (**Fig.4F)**. The right, injured, eye (OD), showed a significant PhNR amplitude alteration between D1 and D7 (p=0.0002), with D1 amplitudes being decreased as compared to elevated D7 amplitudes. Also of note, OD showed a decrease in amplitude at D1 (p=0.0225) as compared to OS. OD D7 PhNR amplitudes were found to significantly overshoot OD Baseline amplitudes (p=0.0225). In the injured group PhNR amplitude significant changes were depicted for the following: OD D1 vs OS D7 (p<0.0001), OD Baseline vs OS D7 (p=0.0012), OS Baseline vs OS D7 (p=0.0008), OS Baseline vs OD D7 (p=0.0094).

**Fig. 4:**
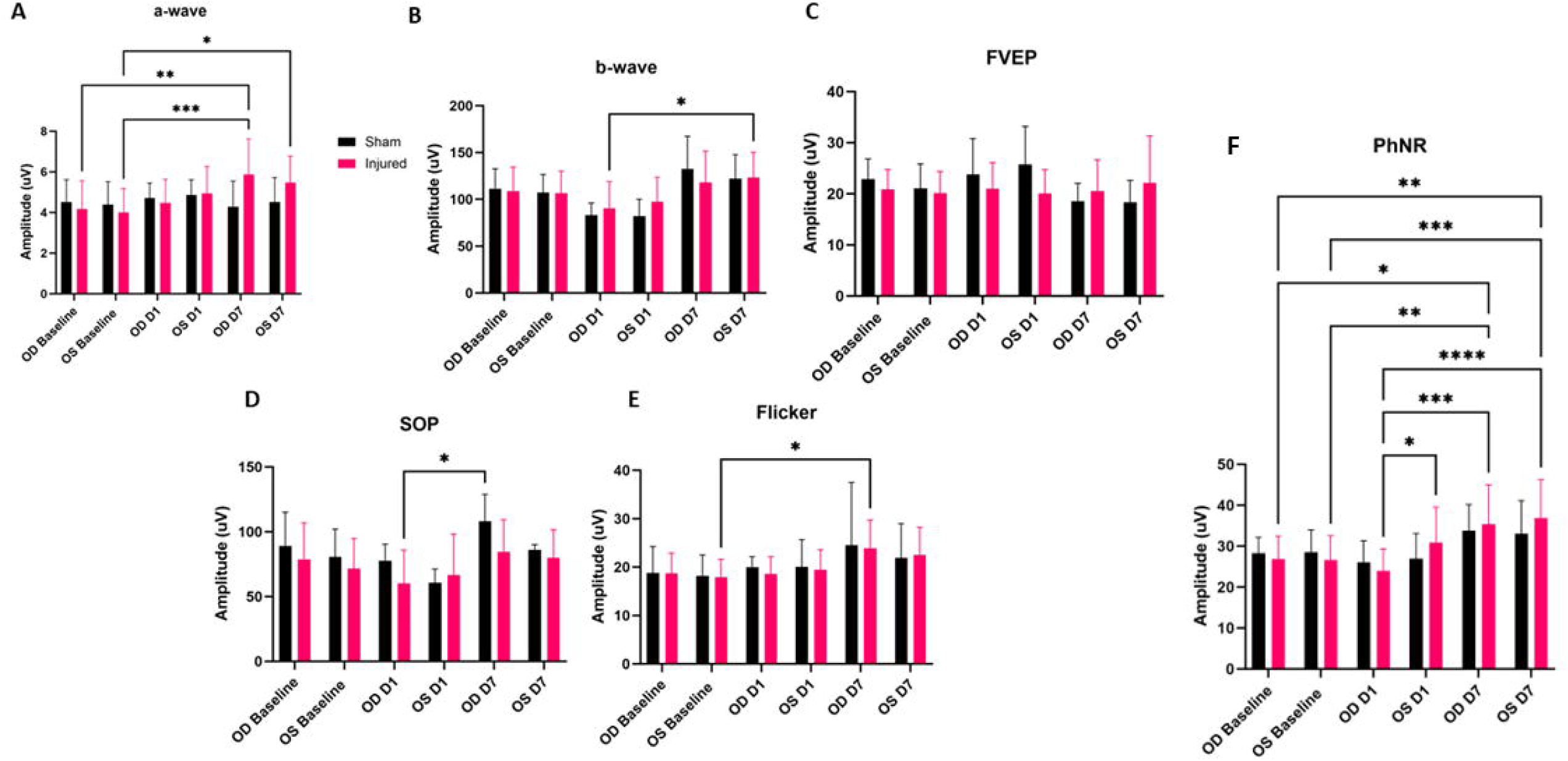
Average ERG amplitudes between injured and experimental groups. ffERG a-wave amplitudes with significant amplitude differences noted between OD Baseline and OD D7 (p=0.0051), OS Baseline and OD D7 (p=0.0010), and OS Baseline and OS D7 (p=0.0294) (A). ffERG amplitudes of OD D1 compared to OS D7 were determined to be statistically significant (B). No statistically relevant differences were noted with regard to FVEP amplitudes (C). SOP amplitudes showed significant alteration between OD D1 and OD D7 (p=0.0251) (D). Flicker amplitudes depicted significant changes between OS Baseline and OD D7 (p=0.0125) (E). Significant differences were noted for the injured animal PhNR amplitudes between OD D1 and OD D7 (p=0.0002), OD D1 and OS D7 (p<0.0001), OD D1 and OS D1 (p=0.0225), OS Baseline and OS D7 (p=0.0008), OS Baseline and OD D7 (p=0.0094), OD Baseline and OS D7 (p=0.0012), and OD Baseline and OD D7 (p=0.0129) (F).

A significant rise in amplitude was detected between OD D7 and OD Baseline a-wave amplitude (p=0.0051). Additionally, we observed a-wave amplitude statistical significances between OS Baseline and OD D7 (p=0.0010) and OS Baseline and OS D7 (p=0.0294). The b-wave showed one statistically significant amplitude increase between OD D1 and OS D7 (p=0.0153). The SOPs had one detectable significant alteration between the injured OD at D1 as compared to the right eye sham control at D7 (p=0.0251). The flicker waveform depicted one statistically significant rise in of amplitude between OS Baseline and OD D7 (p=0.0125). We did not observe any statistically significant alterations in the fVEP waveforms between eyes or between groups. Additionally, we did not observe any significant changes to peak latency for any of the electrophysiological waveforms.

### Histology and imaging

#### Immunofluorescent staining

Fluorescent labeling of the retina in injured eyes revealed increased GFAP labeling at D7 (**Fig. 5C,H**). This suggests the presence of stressed conditions within the retina. Increased GFAP labeling was observed to extend from the ganglion cell layer into the photoreceptor layer, which indicated Müller cells were becoming hypertrophic. This observation was made in eight of the nine analyzed retinal samples. GFAP labeling in injured D1 retinas appeared to be thickening at the RGC layer, but retinas did not appear to have increased Müller glial extensions (**Fig. 5A,F**). This was observed in two out of three retinas analyzed with the third showing potential thickening but was not as widespread as the other retinas. DAPI and β-tubulin did not show a significant alteration between injured eye, contralateral eye, or sham eyes.

**Fig. 5:**
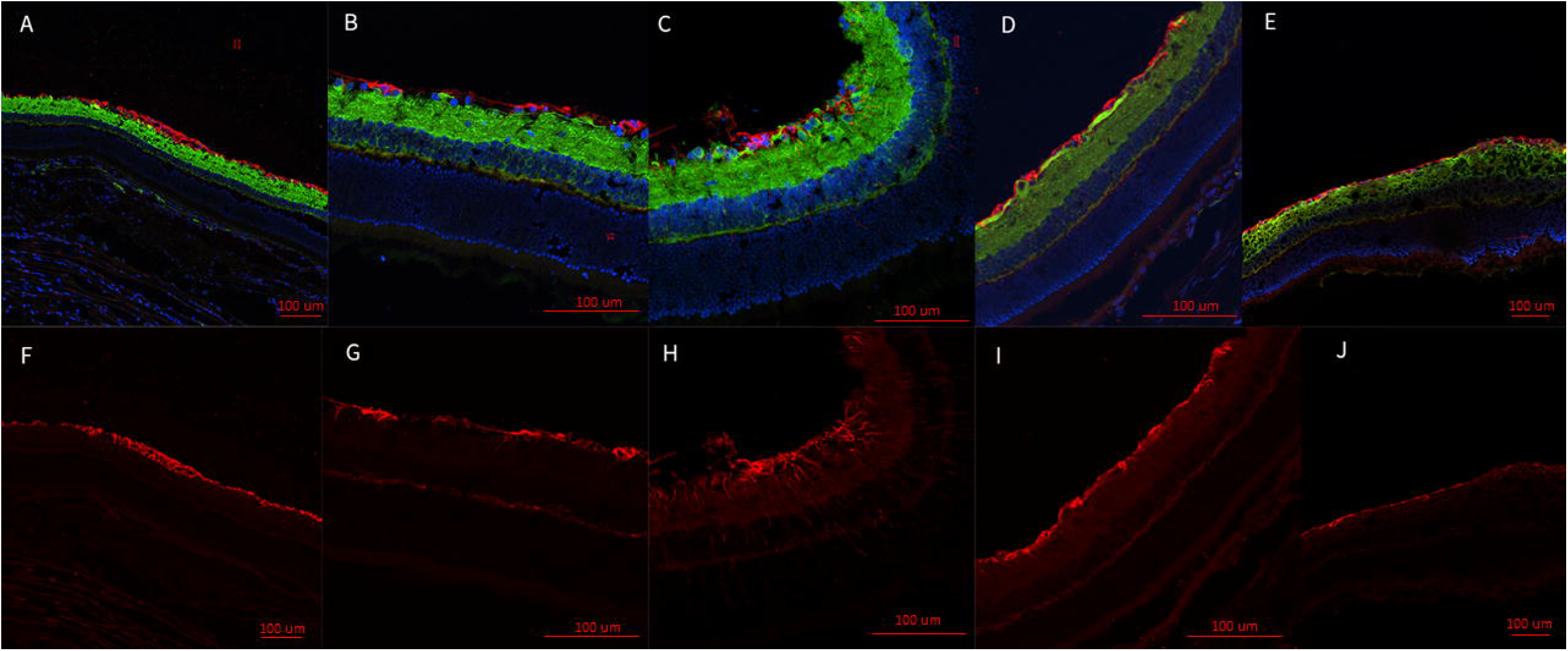
Representative epifluorescence retinal images. Retinas are labeled with the nuclear marker, DAPI, and immunolabeled for the axon marker, β-tubulin (green), and the glial marker, GFAP (red) (A-E). GFAP marked channels (F-J). Injured eye D1 (A,F); contralateral eye (B,G); injured eye D7 (C,H); contralateral eye (D, I); sham (E,J). GFAP appears to be thicker at the retinal ganglion cell (RGC) layer of D1 injured retinas (A,F). The injured eye at D7 was observed to develop increased hypertrophic activity of the Müller glial cells as observed by the increased GFAP presence and the extensions of the Müller glial cells from the RGC bodies through the retina to the photoreceptors (C, H). These extensions were not observed in the contralateral eye at D1 (B,G) or D7 (D, I). Nor were they represented in the sham (E,J), or the injured eye at D1 (A,F).

D7 optic nerve immunohistochemical analysis found decreased labeling intensity β-tubulin in injured nerves (n=3) relative to sham (n=1) (**Fig. 6D**). Changes were observed bilaterally, though not symmetrically, suggesting that the biomechanical mechanism of injury employed must transmit stresses through the chiasm. Biomechanical stress transmission through the chiasm was therefore investigated and directly observed in a cadaver animal following craniotomy.

**Fig. 6:**
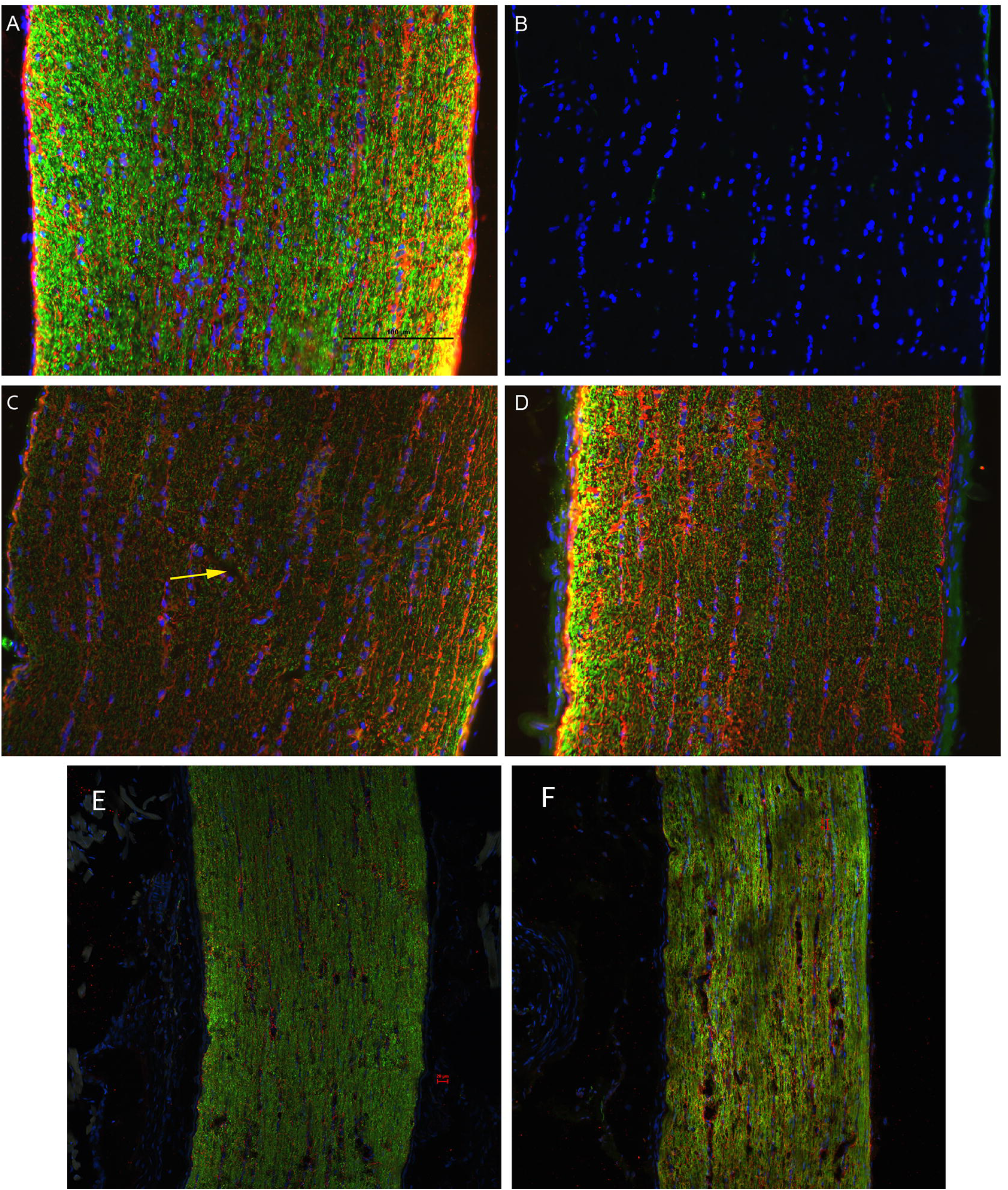
Representative epifluorescence micrographs of longitudinal sections of optic nerves. Optic nerves are labeled with the nuclear marker, DAPI, and immunolabeled for the axon marker, β-tubulin (green), and the glial marker, GFAP (red): uninjured (A); uninjured, no primary antibody (B); left (C) and right (D) optic nerves of D7 injured animal (n=3); left (E) and right (F) optic nerves of D1 injured animals (n=3). In the control (A), β-tubulin is highly punctate with marginally higher intensity near the surface of the nerve; GFAP is most prevalent in localized gaps between β-tubulin and slightly brighter near the surface. (C) In the contralateral ON, β-tubulin has a lower intensity throughout and is disrupted across a diagonal portion of the ON (yellow arrow); GFAP also has a significantly lower intensity across the ON. (D) In the ipsilateral eye, β-tubulin labeling was lower intensity and more diffuse in most of the ON but with large, high-intensity puncta at left; larger bundles of GFAP were colocalized with axial columns of DAPI, but also showed some radial bundling. β-tubulin presence in the injured eye at D1 (F) appears to have a lower intensity as compared to the contralateral eye (E).

#### TUNEL

TUNEL+ cells were detected in the retina and optic nerve of injured animals in both OD and OS (**Fig. 7**). Generally, retinas obtained from animals who received the injury displayed similar counts of TUNEL+ cells in both eyes at each timepoint. Both OD and OS retinas appeared to feature a rise in TUNEL+ cells by D7 (**Fig. 7 A-D,K**). Optic nerve TUNEL staining showed an increased amount of TUNEL+ cells detected in OD samples at both D1 and D7 as compared to the contralateral eye; however, there were no statistically significant differences detected (**Fig. 7 F-J,L**). Retinal and nerve multiple comparisons were made to determine statistical relevance of cell death counts as detailed in **Table 1**.

**Fig. 7:**
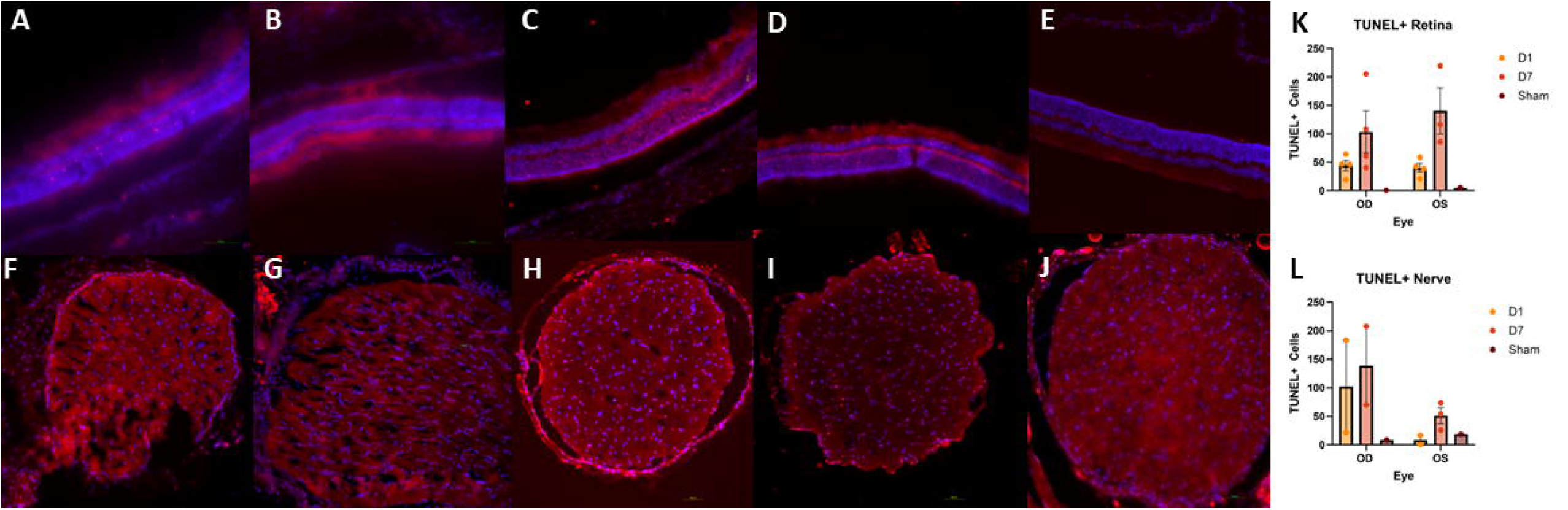
Representative TUNEL micrographs of retina (A-E) and optic nerves (F-J). Injured OD D1 (A,F), Contralateral OS D1 (B,G), Injured OD D7 (C,H), Contralateral OS D7 (D,I), Sham (E,J). Some TUNEL positive (TUNEL+) cells were detected 24 hours after injury (D1) with an increase in TUNEL+ cells in both retina and optic nerve occurring by 7 days after injury (D7) (K,L). TUNEL+ cell counts were higher in OD optic nerves as compared to OS optic nerves at both D1 and D7 (L).

**Table 1.** TUNEL Tukey Multiple Comparisons for the Retina and Optic Nerve.

| TUNEL | Retina | Nerve |
| --- | --- | --- |
| Comparisons | p-value | p-value |
| OD :D1 vs. OD :D7 | 0.5731 | 0.9921 |
| OD :D1 vs. OD :Sham | 0.9667 | 0.8593 |
| OD :D1 vs. OS:D1 | >0.9999 | 0.7493 |
| OD :D1 vs. OS:D7 | 0.2006 | 0.9527 |
| OD :D1 vs. OS:Sham | 0.9784 | 0.9031 |
| OD :D7 vs. OD :Sham | 0.4858 | 0.6592 |
| OD :D7 vs. OS:D1 | 0.5108 | 0.4959 |
| OD :D7 vs. OS:D7 | 0.9204 | 0.7327 |
| OD :D7 vs. OS:Sham | 0.5271 | 0.718 |
| OD :Sham vs. OS:D1 | 0.9778 | >0.9999 |
| OD :Sham vs. OS:D7 | 0.2322 | 0.9917 |
| OD :Sham vs. OS:Sham | >0.9999 | >0.9999 |
| OS:D1 vs. OS:D7 | 0.1723 | 0.9776 |
| OS:D1 vs. OS:Sham | 0.9865 | >0.9999 |
| OS:D7 vs. OS:Sham | 0.2573 | 0.9976 |

Multiple comparisons were made between timepoints (Baseline, D1, and D7) and between groups (injured and sham). Comparisons were also made between eyes (OD injured and OS contralateral).

## Discussion

Based on our findings and clinical considerations, we believe that the PhNR of the LA ffERG may be the most beneficial test for early diagnosis of TON since we observed statistically significant alterations in amplitudes between the injured eye and contralateral eye 24 hours after injury inducement (D1), and we also observed a significant alteration in PhNR amplitude between D1 and D7 (**Fig. 4**) (8,19). Our retinal immunofluorescent staining supports these findings with our observations of increased GFAP staining at D1, via the thickening of Müller glial cell bodies in the RGC layer of the retina at D1 and then the increased hypertrophic activity of the Müller glial cells observed in the retinas of animals at D7 (**Fig.5**). Epifluorescence indicated significant changes in both β-tubulin and GFAP labeling within the injured optic nerve (**Fig. 6**). Reduced β-tubulin labeling suggests a decrease in the number of microtubules within each axon and/or fewer axons. Furthermore, we observed an increase in TUNEL+ cells in injured retinas and optic nerves, supporting our model of torsion-induced TON. Combined with diminished PhNRs, these findings support the hypothesis that our model is causing a loss of activity in the ganglion cells, which is indicative optic neuropathy.

Together, these findings provide novel insights into the secondary neurodegenerative mechanisms leading to vision loss from TON. The developed animal model represents an excellent platform for studying TON, developing diagnostics, and evaluating candidate treatments for safety and efficacy. This model has tremendous potential for evaluation of both pre- and post-traumatic therapies.

ffERGs are a series of visual stimuli-electrophysiologic response waveforms which serve to analyze the functional output of the various cell types within the visual system. Light adapted ERGs require no dark adaptation time, which makes them relatively faster for real world injury applications. The different cell types within the retina can be observed through the inducement of various light stimulating protocols. Generally, the a-wave of the LA ffERG consists of the photoreceptor response. The a-wave is the first waveform of the ffERG and arises due to the hyperpolarization of the photoreceptor membranes during phototransduction (20–23). The b-wave is the positive component of the ffERG signal and reflects the depolarization of the bipolar cells (20–23).The OPs are observed in the ffERG as the oscillations on the ascending limb of the b-wave, and in rodents the OPs can also be on the descending limb as well (22–24). These signals are generated by the inhibitory effect of the amacrine cells. The amacrine cells release γ-aminobutyric acid (GABA) and glycine, which act as inhibitory neurotransmitters, and create the oscillatory features observed (22,23). The PhNR is the negative deflection observed after the b-wave, and is representative of retinal ganglion cell (RGC) and amacrine cell activity (25–27). The VEP is a waveform which can assess the post-retinal function by characterizing the electrical output of the visual cortex (28). The light-adapted flicker ERG at 20Hz is known to isolate the cone function in rat retinas (29). With analysis of each of these waveforms we can determine which cell types are afflicted after injury and find a diagnostic criterion for TON.

During our ffERG analysis we noticed the sham animals were displaying rising signals similar to our left (contralateral) eye (OS) recordings. Repetitive ketamine dosing may have impacted our inner retinal cells as the inner retinal ffERG waveforms such as the PhNR, b-wave, and SOPs displayed increasing signals over time. This phenomenon has been reported in other literature and we believe ketamine has been acting on these inner retinal cells via the mechanisms outlined in **Figure 8** (30–34). Briefly, after TBI, glutamate levels rise, and NMDA receptor function increases, and ketamine has been reported to block both glutamate and NMDA receptors. The highest concentration of NMDA receptors are found in the inner retinal cells, which are also the cell types dominating the PhNR, b-wave, and SOPs (33). Due to this, we believe the back-to-back ketamine doses at injury and then 24 hours later for D1 ERG may be depressing the inner retinal ERG dependent waveforms. This would be why our sham animals are following the same trend of our contralateral controls. Our study design inherently controls for this by ensuring all animals have the same number and timing of anesthetic events. In addition, we normalized the electroretinogram data against baseline measurements, but there were not any significant changes between the non-normalized and normalized datasets, indicating changes observed were significant to the model and not the result of miscellaneous factors.

**Fig. 8:**
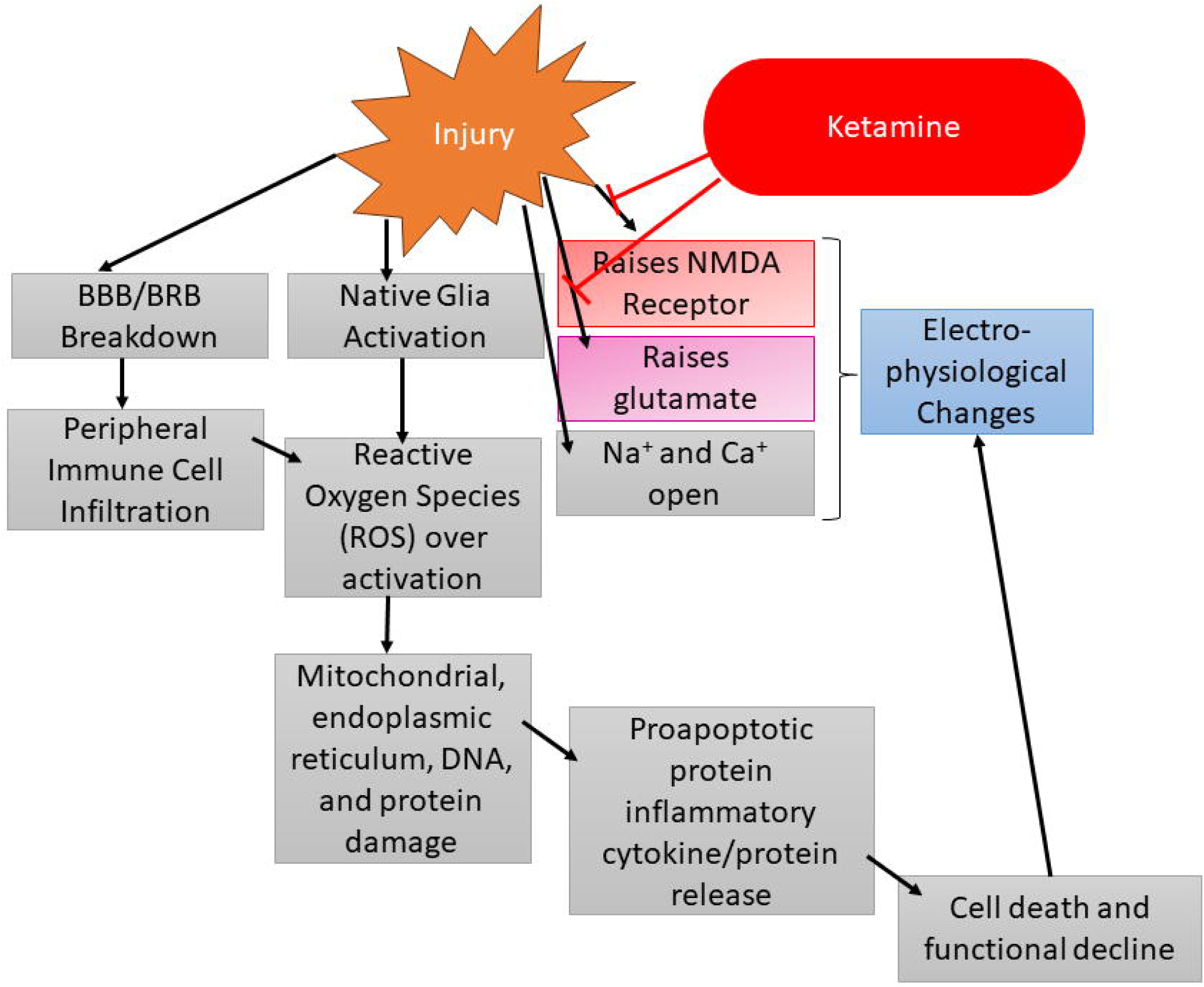
Potential flow chart of the injury cascade of events following TBI/TON inducement. Importantly, the injury event increases the activity of the NMDA receptor and elevates glutamate levels. Both of these factors are then attenuated by ketamine, which can lead to electrophysiological changes.

The observation of ketamine’s effect after repetitive dosing poses an interesting theory as to whether it can be used as a neuroprotective therapeutic, especially since it can target inner retinal cells via the suppression of the NMDA receptor. One rodent study investigating neuroprotection after light induced damage reported ketamine/xylazine as a potential pretreatment neuroprotectant (32). In the future we plan to elucidate therapeutic strategies for retinal cell preservation. Alterations of the PhNR amplitude indicate inner retinal cell damage, so therapeutics capable of preservation of this area of the retina may prove to be the most beneficial in minimizing cell and vision loss after injury. Other potential therapeutic strategies could target the anti-inflammatory aspect of TON. The inflammatory response that happens in the days following the primary injury may be just as harmful if not more damaging than the initial injury event. Being able to suppress this inflammatory response may help to prevent the subsequent damage characteristic of TON. Potential therapeutics for this type of therapeutic intervention include ibudilast, TUDCA, cyclosporine, erythropoietin, dopamine, and vitamins such as C and E (1,35–41).

This study has potential limitations. One such limitation was the utilization of only one animal sex for each portion of the study: animals were either all female or all male but did not include subjects from both sexes for each sub experiment. Sex and weight differences were not accounted for between the sub experiments, nor were they evaluated, thus presenting an inherent limitation. Due to timing the main experiment utilized male rodents as their hormonal levels did not change throughout the month, resulting in more consistent ffERGs. In future work, both sexes of animals should be evaluated for diagnostic and therapeutic intervention strategies. Generally, men are reported to have the highest incidence of TBI; however, women suffering TBI have a greater risk of death as compared to male counterparts (42). This indicates the biomechanical mechanisms governing TBI and potentially TON may differ with regard to sex. Another limitation to this study is we utilized LA ffERGs, which target more cone receptors and rodents are known to be a rod dominated species (43). We may have observed larger deficits in ffERG amplitudes had we utilized dark adapted ffERGs. LA ffERGs were chosen as they are quicker to conduct in a clinical setting since they avoid dark adaptation time. Finally, the mechanism of TON induction used in this study was designed to provide a repeatable mechanical stimulus rather than mimic what might be considered a more typical TON-inducing injury. This decision was taken to maximize experimental precision and thereby reduce the number of animals required to achieve statistically significant results when evaluating candidate therapeutics. In reality, TON may be caused by many unique insults; however, the end result of RGC death and irreversible visual dysfunction is thought to be quite similar to what we observed in our model. Another limitation to the study was due to our small animal numbers allocated to immunohistochemical staining, we relied on descriptive results and analysis of tissue structures. While some optical coherence tomography (OCT) work was performed, no significant changes were noted, presumably due to the short-term follow-up period (7 days). Future studies will also utilize OCT to evaluate potential thickness changes in the retina after TITON. In addition, through our TUNEL analysis, we were able to determine general cell death in both the retina and optic nerve, but in future work we plan to analyze retinal flat mounts in order to further elucidate specific cells dying after TITON. In addition, we plan to analyze different immunohistochemical staining techniques, such as Nestin and S100β, to determine which glial cells are maximally activated after TITON.

### Conclusions

We have shown that our torsion-induced TON animal model can produce a diagnosable functional change in ffERGs via analysis of the PhNR. This functional change correlates with stressed conditions in the retina as evidenced by our tissue analysis. This is the first step in understanding how the retina and optic nerve are afflicted after TON insult. Future studies will determine therapeutic intervention strategies capable of electrophysiological function in the retinal neuronal circuits. Through our therapeutic intervention we hope to elucidate the cellular mechanisms responsible for the characteristic visual system deterioration of TON. By selecting therapeutics which target different suspected pathways of secondary neurodegeneration we hope to determine how TON progresses and at which time points therapeutic intervention would be the most efficacious (1).

### Transparency, Rigor, and Reproducibility

All anesthetic events were controlled by utilizing the same dosing schedule and quantities between animals and groups. In addition, ffERG amplitude differences were controlled by comparing animals of the same sex and by collecting all visual electrophysiological data during the same time of day. This set of controls negates sex-based differences and then also ensures circadian rhythm effects have been minimized. All animals used were withing the same weight range, minimizing animal-to-animal differences. Both sham animals and contralateral eyes (OS) were used as controls. Optic nerves cross with partial decussation at the chiasm, but nerve fibers never coalesce to form new, unique fibers. We hypothesized there may be an intrachiasmal transference between the nerves due to our injury event. As such, we utilized sham animals as an additional control parameter. The injury device is controlled and allows for a higher rate of reproducibility as the motorized system executes the torsional motion. This motion is preprogrammed in terms of degrees, velocity, and torque. This allows the user to ensure the parameters of the injury event are the same between animals.

## Acknowledgements

We would like to thank The Ohio State University laboratory and animal resources (ULAR), Dr. Stacey Meeker, Dr. Colleen Cebulla. In addition, we would like to thank Reilly undergraduate lab members: Stephanie Small, Michelle Mosko, Emma Lally, Vidhya Kannan, Sam Duckworth, and Eve Howard. We would also like to thank Dr. Wade Rich and Elizabeth Urbanski.

## Supporting Information

**S1 File. Data tables.** Data tables required to reproduce all statistical analyses. Each row denotes the independent variable. Each column represents data from a single animal.

## References

1. Ryan AK, Rich W, Reilly MA. Oxidative stress in the brain and retina after traumatic injury. Front Neurosci [Internet]. 2023;17. Available from: https://www.frontiersin.org/journals/neuroscience/articles/10.3389/fnins.2023.102115 2

2. Chan JW. Optic Nerve Disorders: Diagnosis and Management [Internet]. 1st ed. New York, NY: Springer; 2008. 284 p. Available from: 10.1007/978-0-387-68979-1

3. Sarkies N. Traumatic optic neuropathy. Eye. 2004 Nov 1;18(11):1122–5.

4. Prevention C for DC and. Report to congress: Traumatic brain injury in the United States. 2016; Available from: https://www.cdc.gov/traumaticbraininjury/pubs/tbi_report_to_congress.html

5. Scott R. The injured eye. Philos Trans R Soc B Biol Sci. 2011 Jan 27;366(1562):251–60.

6. Sherwood D, Sponsel WE, Lund BJ, Gray W, Watson R, Groth SL, et al. Anatomical Manifestations of Primary Blast Ocular Trauma Observed in a Postmortem Porcine Model. Invest Ophthalmol Vis Sci. 2014;55(2):1124–32.

7. Steinsapir KD, Goldberg RA. Traumatic optic neuropathy. Surv Ophthalmol. 1994 Jun;38(6):487–518.

8. Blanch RJ, Joseph IJ, Cockerham K. Traumatic optic neuropathy management: a systematic review. Eye. 2024 Aug 1;38(12):2312–8.

9. Steinsapir KD, Goldberg RA. Traumatic optic neuropathy: an evolving understanding. (1879-1891 (Electronic)).

10. Ian Roberts, Yates D, Sandercock P, Farrell B, Wasserberg J, Lomas G, et al. Effect of intravenous corticosteroids on death within 14 days in 10008 adults with clinically significant head injury (MRC CRASH trial): randomised placebo-controlled trial. Lancet. 2004 Oct;364(9442):1321–8.

11. Tse BC, Dvoriantchikova G, Tao W, Gallo RA, Lee JY, Pappas S, et al. Tumor Necrosis Factor Inhibition in the Acute Management of Traumatic Optic Neuropathy. Invest Ophthalmol Vis Sci. 2018 Jun 1;59(7):2905–12.

12. Khan RS, Ross AG, Aravand P, Dine K, Selzer EB, Shindler KS. RGC and Vision Loss From Traumatic Optic Neuropathy Induced by Repetitive Closed Head Trauma Is Dependent on Timing and Force of Impact. Transl Vis Sci Technol. 2021 Jan;10(1):8.

13. Burke EG, Cansler SM, Evanson NK. Indirect traumatic optic neuropathy: modeling optic nerve injury in the context of closed head trauma. Neural Regen Res [Internet]. 2019;14(4). Available from: https://journals.lww.com/nrronline/fulltext/2019/14040/indirect_traumatic_optic_neuropathymodeling.9.aspx

14. Bricker-Anthony C, Hines-Beard J, Rex TS. Molecular changes and vision loss in a mouse model of closed-globe blast trauma. Invest Ophthalmol Vis Sci. 2014 Jul 3;55(8):4853–62.

15. Li Y, Singman E, McCulley T, Wu C, Daphalapurkar N. The Biomechanics of Indirect Traumatic Optic Neuropathy Using a Computational Head Model With a Biofidelic Orbit. Front Neurol. 2020;11:346.

16. Galgano M, Toshkezi G, Qiu X, Russell T, Chin L, Zhao LR. Traumatic Brain Injury: Current Treatment Strategies and Future Endeavors. Cell Transplant. 2017 Jul;26(7):1118–30.

17. You Y, Klistorner A, Thie J, Graham SL. Improving reproducibility of VEP recording in rats: electrodes, stimulus source and peak analysis. Doc Ophthalmol Adv Ophthalmol. 2011 Oct;123(2):109–19.

18. Heisler-Taylor T, Kim B, Reese AY, Hamadmad S, Kusibati R, Fischer AJ, et al. A new multichannel method quantitating TUNEL in detached photoreceptor nuclei. Exp Eye Res. 2018 Nov;176:121–9.

19. Dhillon A, Ahmad MSZ, Breeze J, Blanch RJ. Prolonged deployed hospital care in the management of military eye injuries. Eye Lond Engl. 2020 Nov;34(11):2106–11.

20. Perlman I. University of Utah Health Sciences Center. In: Webvision: The Organization of the Retina and Visual System [Internet] [Internet]. Salt Lake City, UT: University of Utah Health Sciences Center; 2001. Available from: https://www.ncbi.nlm.nih.gov/books/NBK11554/

21. Reichel E, Klein K. Retinal Electrophysiology. In: Ophthalmology [Internet]. 5th ed. Elsevier; 2019. Available from: https://www.clinicalkey.com/#!/content/book/3-s2.0-B9780323528191002875

22. Audo I, Robson AG, Holder GE, Moore AT. The negative ERG: clinical phenotypes and disease mechanisms of inner retinal dysfunction. Surv Ophthalmol. 2008 Feb;53(1):16–40.

23. Mallery R, Mackay D, Prasad S. Ocular Functional and Structural Investigations. In: Bradley and Daroff’s Neurology in Clinical Practice [Internet]. 8th ed. Elsevier; 2022. p. 601–13. Available from: https://www.clinicalkey.com/#!/content/book/3-s2.0-B9780323642613000437?scrollTo=#23hl0000474

24. Rosolen SG, Rigaudière F, LeGargasson JF, Chalier C, Rufiange M, Racine J, et al. Comparing the photopic ERG i-wave in different species. Vet Ophthalmol. 2004 Jun;7(3):189–92.

25. Li B, Barnes GE, Holt WF. The decline of the photopic negative response (PhNR) in the rat after optic nerve transection. Doc Ophthalmol Adv Ophthalmol. 2005 Jul;111(1):23–31.

26. Jnawali A, Lin X, Patel NB, Frishman LJ, Ostrin LA. Retinal ganglion cell ablation in guinea pigs. Exp Eye Res. 2021 Jan;202:108339.

27. Miura G, Wang MH, Ivers KM, Frishman LJ. Retinal pathway origins of the pattern ERG of the mouse. Exp Eye Res. 2009 Jun 15;89(1):49–62.

28. Ridder WH, Nusinowitz S. The visual evoked potential in the mouse—Origins and response characteristics. Vision Res. 2006;46(6):902–13.

29. Goto Y, Yasuda T, Tobimatsu S, Kato M. 20-Hz flicker stimulus can isolate the cone function in rat retina. Ophthalmic Res. 1998;30(6):368–73.

30. Choh V, Gurdita A, Tan B, Feng Y, Bizheva K, McCulloch DL, et al. Isoflurane and ketamine:xylazine differentially affect intraocular pressure-associated scotopic threshold responses in Sprague-Dawley rats. Doc Ophthalmol Adv Ophthalmol. 2017 Oct;135(2):121–32.

31. Prando S, Carneiro C de G, Otsuki DA, Sapienza MT. Effects of ketamine/xylazine and isoflurane on rat brain glucose metabolism measured by (18) F-fluorodeoxyglucose-positron emission tomography. Eur J Neurosci. 2019 Jan;49(1):51–61.

32. Arango-Gonzalez B, Schatz A, Bolz S, Eslava-Schmalbach J, Willmann G, Zhour A, et al. Correction: Effects of Combined Ketamine/Xylazine Anesthesia on Light Induced Retinal Degeneration in Rats. PLOS ONE. 2012 Aug 6;7(8):10.1371/annotation/d52a610e-6a56-4158-90bf-45e37f053567.

33. Shen Y, Liu XL, Yang XL. N-methyl-D-aspartate receptors in the retina. Mol Neurobiol. 2006 Dec;34(3):163–79.

34. Deng Y, Thompson BM, Gao X, Hall ED. Temporal relationship of peroxynitrite-induced oxidative damage, calpain-mediated cytoskeletal degradation and neurodegeneration after traumatic brain injury. Exp Neurol. 2007 May;205(1):154–65.

35. Bond WS, Rex TS. Evidence That Erythropoietin Modulates Neuroinflammation through Differential Action on Neurons, Astrocytes, and Microglia. Front Immunol. 2014;5:523.

36. Daruich A, Picard E, Boatright JH, Behar-Cohen F. Review: The bile acids urso- and tauroursodeoxycholic acid as neuroprotective therapies in retinal disease. Mol Vis. 2019;25:610–24.

37. Cueva Vargas JL, Belforte N, Di Polo A. The glial cell modulator ibudilast attenuates neuroinflammation and enhances retinal ganglion cell viability in glaucoma through protein kinase A signaling. Neurobiol Dis. 2016 Sep;93:156–71.

38. Bernardo-Colón A, Vest V, Clark A, Cooper ML, Calkins DJ, Harrison FE, et al. Antioxidants prevent inflammation and preserve the optic projection and visual function in experimental neurotrauma. Cell Death Dis. 2018 Oct 26;9(11):1097.

39. DeJulius CR, Bernardo-Colón A, Naguib S, Backstrom JR, Kavanaugh T, Gupta MK, et al. Microsphere antioxidant and sustained erythropoietin-R76E release functions cooperate to reduce traumatic optic neuropathy. J Control Release Off J Control Release Soc. 2021 Jan 10;329:762–73.

40. Lan YL, Li S, Lou JC, Ma XC, Zhang B. The potential roles of dopamine in traumatic brain injury: a preclinical and clinical update. Am J Transl Res. 2019;11(5):2616–31.

41. Lendvai-Emmert D, Magyar-Sumegi ZD, Hegedus E, Szarka N, Fazekas B, Amrein K, et al. Mild traumatic brain injury-induced persistent blood–brain barrier disruption is prevented by cyclosporine A treatment in hypertension. Front Neurol [Internet]. 2023;14. Available from: https://www.frontiersin.org/journals/neurology/articles/10.3389/fneur.2023.1252796

42. Munivenkatappa A, Agrawal A, Shukla DP, Kumaraswamy D, Devi BI. Traumatic brain injury: Does gender influence outcomes? Int J Crit Illn Inj Sci. 2016 Jun;6(2):70–3.

43. Galindo-Romero C, Norte-Muñoz M, Gallego-Ortega A, Rodríguez-Ramírez KT, Lucas-Ruiz F, González-Riquelme MJ, et al. The retina of the lab rat: focus on retinal ganglion cells and photoreceptors. Front Neuroanat [Internet]. 2022;16. Available from: https://www.frontiersin.org/journals/neuroanatomy/articles/10.3389/fnana.2022.9948 90

